# The possibility of anesthesia stimulated by a train of current pulses

**DOI:** 10.1101/2022.10.13.512150

**Authors:** Mikhail N. Shneider, Mikhail Pekker

## Abstract

This paper considers a simple theoretical model of blocking the passage of signals (action potentials) from sensory neurons and thereby effecting anesthesia without the use of anesthetics as a result of a sequence of unipolar current pulses generated by an external source. The proposed model allows the selection of parameters and the required frequency of the repetition of current pulses for the possible implementation of anesthesia depending on the electrical characteristics of the skin and the conductivity of the saline solution in which the myelinated nerve fibers are located.

## 1. Introduction

Acupuncture as a means to relieve pain has been known since the third millennium BC and has remained essentially the same to the present day [1-5]. One of the modern innovations to the traditional acupuncture technique, however, is the use of weak alternating current sources connected to needles (electroacupuncture). This has been shown to significantly improve the effect of anesthesia [6-8]. It should be noted that [9 (PRE2014)] considered the possibility of anesthesia associated with the redistribution of transmembrane ion channels in the membrane of nerve cells in the acoustic field of an ultrasound source or in the field of stimulated longitudinal ultrasonic oscillations resulting from the interaction of a charged membrane with microwaves.

The research in [10-18] shows the effectiveness of restoring control over the lower extremities during the spinal stimulation of the lumbosacral areas of the spinal cord below the site of spinal cord injury using non-invasive electrodes placed directly on the skin above the lower spinal cord. Additionally, a simple theoretical model of the excitation of the action potentials of multiple motor pools by stimulating current pulses over the lumbosacral regions of the spinal cord was considered in [19].

This paper shows that a similar approach makes it possible in principle to block the passage of signals from sensory neurons with the help of external electrodes and thus to perform anesthesia without the use of anesthetics. An example of such an approach would be finger anesthesia (Fig. 1(A)), in which signals from receptors on the fingertips, such as heat receptors, are blocked. This approach, which allows the experimental testing of the theory, entails a dielectric cylinder with ring electrodes in direct contact with the skin, worn, for example, on a finger. Let us consider a model problem in which a nerve fiber (myelinated axon) of radius *a* is located on the axis of a hollow dielectric cylinder of radius *R*_0_ (*a* ≪ *R*_0_) and filled with a conductive liquid (saline) with specific conductivity *σ* (Fig. 1(B)). The currents inside the cylinder, generated by ring electrodes, are fed with short unipolar current pulses, as shown in Fig. 2. Section II of this paper shows that the problem of generating currents in saline inside a dielectric cylinder can be considered within the framework of a stationary continuity equation, similar to that used by us, the authors, earlier in stimulation theory [19]. In Section III, the corresponding distributions of the potential and currents inside a dielectric cylinder with built-in ring electrodes are obtained. Section IV shows that the potential distribution found in section III can lead to the blocking of action potential propagation in the myelinated nerve fiber, as shown in Fig. 1B.

**Figure 1.**
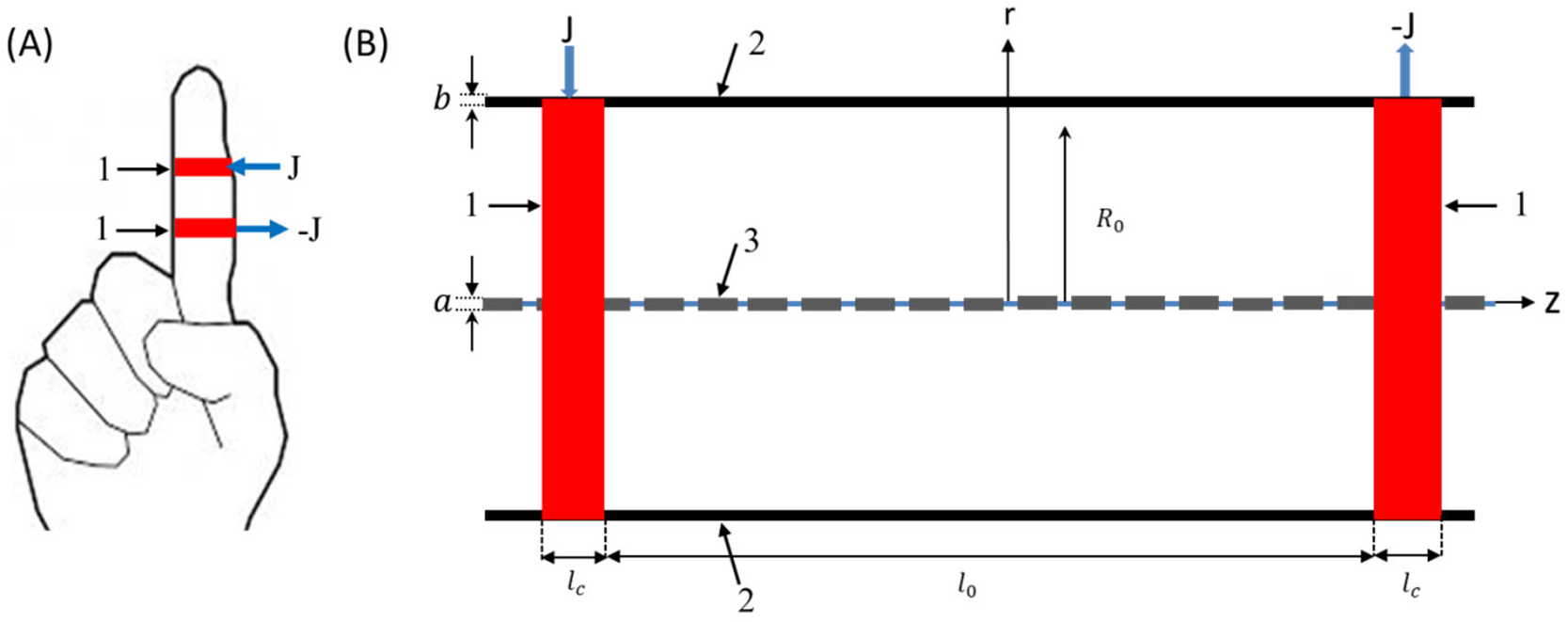
(A) Configuration of external ring electrodes and currents J in the considered example of finger anesthesia; (B) Corresponding model scheme: 1. Electrodes built-in in dielectric cylinder; 2. Radius and thickness of the cylinder are *R*_0_ and *b* correspondently, (*b* ≪ *R*_0_). The cylinder is filled with physiological saline with conductivity σ. 3. Myelinated nerve fiber (axon) of diameter *a* (*a* ≪ *R*_0_) placed on the axis the cylinder. Wide arrows show currents at electrodes 1. *l*_c_, *l*_0_ are the width of the electrodes and the distance between them, respectively.

**Figure 2.**
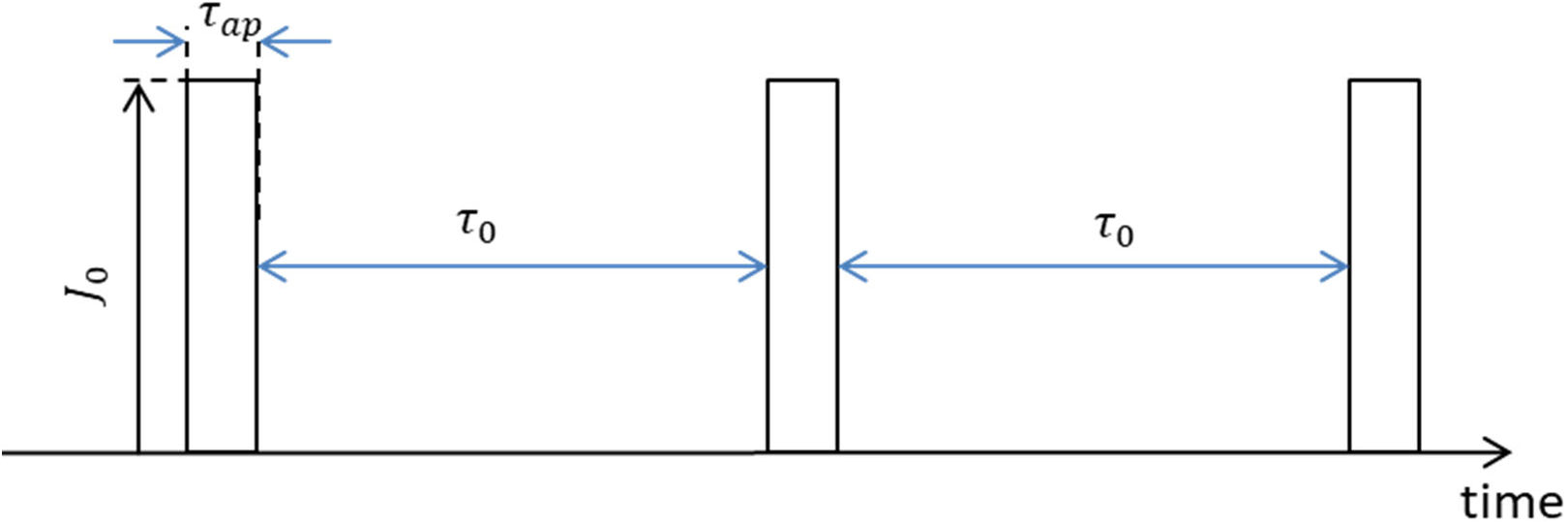
Periodical unipolar current pulses applied to the electrodes. *τ*_ap_, *τ*_0_, and *J*_0_ are the current pulse duration, duty cycle, and amplitude, respectively. Since the duration of the current pulse that blocks the propagation of the action potential must be no less than the time of initiation of the action potential *τ*_1_∼1 ms [20], it is natural to assume that *τ*_ap_∼*τ*_1_. Since the relaxation time of the action potential *τ*_relax_ ∼10–15 ms [20], the duty cycle of the current pulses *τ*_0_∼*τ*_relax_.

## II. Theoretical model

Saline, the fluid in which the neurons are located in living organisms, is a highly conductive electrolyte, with σ∼1 −3 Ω^−1^*m*^−1^ [20]. Therefore, the relaxation time of the volume charge perturbation in saline (the Maxwell time) is of the order 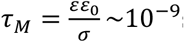, where ε is the dielectric constant of the vacuum, and ε 80 is the relative dielectric permittivity of water, that is, shorter than the characteristic time of the excitation and relaxation of the action potential in the axons by 5–6 orders of magnitude. The size of the non-quasi-neutral region near the membrane of axons is determined by the Debye length 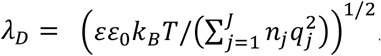. Here, K_B_ is the Boltzmann constant, *T* is the temperature, and *n*_j_ is the densities of the ions with charges *q*_j_. In the case of saline, we can assume that all the ions in the liquid are singly charged and that their total density is of the order *n*_j_ ≈2 ∙10^26^ m^−3^ [22]. At *T* ≈300 *K, λ*_D._ ≈ 0.5 nm, and this value is much smaller than the typical radius of the axon *a* ≈1–8 μm [23]. Therefore, the violation of quasi-neutrality cannot be taken into account, and the potential distribution in the vicinity of a neuron can be found from the equation of the continuity of current:

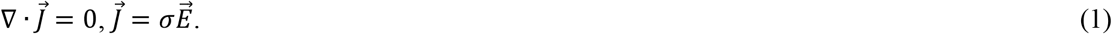

Since the conductivity σ of the electrolyte is uniform and constant in time, and, 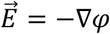 the problem of determining the potential is reduced to the Laplace equation:

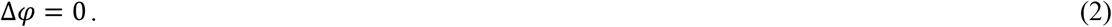

Let us estimate the charging time of the myelinated axon by the currents induced inside the dielectric cylinder (Fig. 1(B)). Since the action potential is generated in unmyelinated sections, nodes of Ranvier, we will consider the charging of the membrane in the nodes of Ranvier, in which the capacitance per unit area 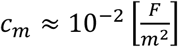and the typical radius *a*∼1.5-8 μm [23].

Following [19], the estimation of the charging time of a cylindrical capacitor of radius *a*, capacitance per unit area *c*_m_, and conductivity σ is:

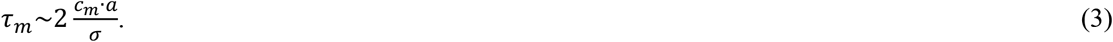

Substituting in (3) the above values *c*_m_, *a*, and σ = 2 Ω^−1^m^−1^, we find that the charging time of the surface of the nodes of Ranvier of the myelinated axon lies in a range from 0.035 to 0.1 μs.

Now let us estimate the charging time for the surface of the outer dielectric cylinder shown in Figure 1(B) with the capacity per unit area

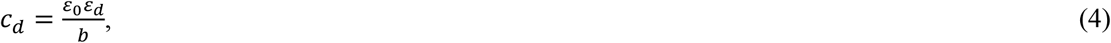

where *b*, ε_d_ are the thickness and relative permittivity of the cylinder wall (Fig. 1(B)). Let us assume that the permittivity of the outer cylinder and its thickness are respectively equal to the permittivity of the ski ε_d_ = ε_skin_ = 12000 [24] and its thickness *b* = *b*_skin_∼0.5–4 mm [25]. Assume, for estimates, *b* = 2 mm. Substituting these values into (4), we get 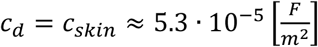. For definiteness, we take the radius of the outer cylinder (finger) *R* = *R*_0_ = 0.01 m. Substituting these values into (3), we find that the charging time of the external cylindrical capacitor is 0.05μs. That is, the charging time of the Ranvier interception membranes in the internal nerve fiber and the external cylindrical capacitor is at least three orders of magnitude less than the duration of the current pulse *t*_ap_ shown in Figure 2. Therefore, during the action of the current pulse, the charge on the fiber membrane and on the surface of the cylinder can be considered constant. Note that the surface of the myelin fiber can be considered as an impenetrable dielectric surface only until the potential on the membrane at the nodes of Ranvier reaches the threshold of the excitation of the action potential. Below we assume that this condition is satisfied.

## III. Potential and current distribution inside the outer dielectric cylinder

Equation (2) in cylindrical coordinates has the form:

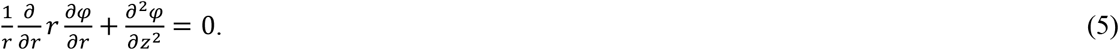

As noted in the previous part of the work, during most of the stimulated current pulse, the charge on the myelin fiber membrane and on the dielectric surface of the outer cylinder can be considered constant. In this case, we can assume that the radial currents on them are equal to zero:

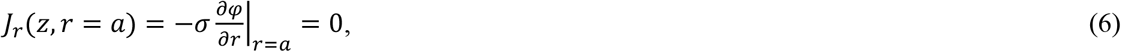

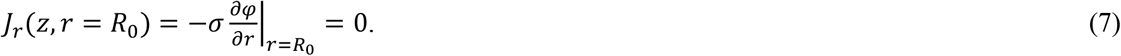

Since *a* ≪ *R*_0_, the influence of the nerve fiber on the distribution of the currents induced by external electrodes inside the dielectric cylinder is insignificant. Therefore, below the presence of myelin fiber will be neglected.

From the symmetry of the problem of current distribution inside a dielectric cylinder with ring electrodes (Fig. 1B) (without myelin fibers on the axis), it follows that the radial current on the axis is:

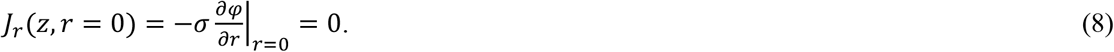

The boundary conditions for the potential φ on the inner surface of the dielectric cylinder have the form:

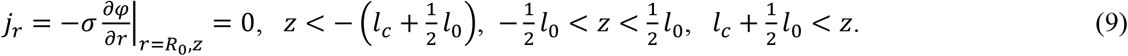

On the ring electrodes, the current is set as follows:

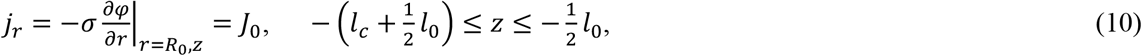

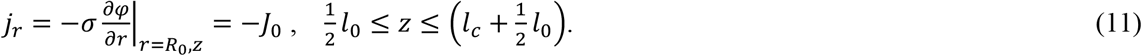

Without loss of generality, the stepwise longitudinal current distribution on the electrodes can be approximated by the sum of the exponents:

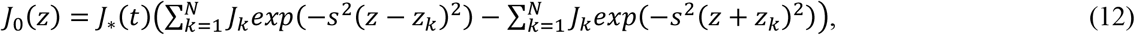

where *J*_∗_(*t*) is a step function corresponding to Figure (2), and *J*_k_, *z*_k_ are constant values. Assuming *J*_k_ = 1, we obtain the distribution of currents and potential inside the dielectric cylinder [26-28]:

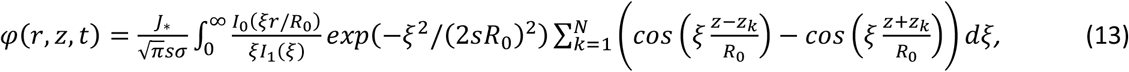

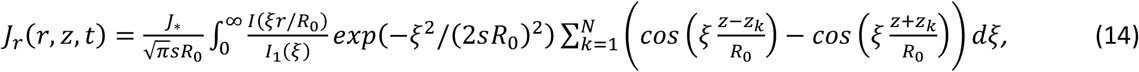

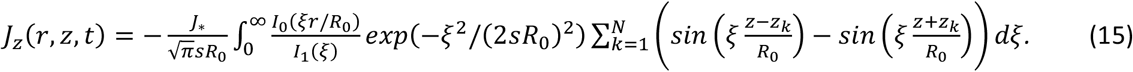

*I*_0_, *I*_1_ in (13)-(15) are modified Bessel functions of the first kind.

Figure 3 shows an example of the calculation results for the selected maximum current on the electrode *J*_0_ = 0.15A, dimensions *R*_0_ = 1 cm, *l*_0_ = 10 cm, *z*_1_ = 5 cm, *z*_2_ = 5.25 cm, *z* = 0.5 cm, *z* = 0.575 cm, *z*_5_ = 6 cm, and *s* = 5 cm^−1^.

**Figure 3.**
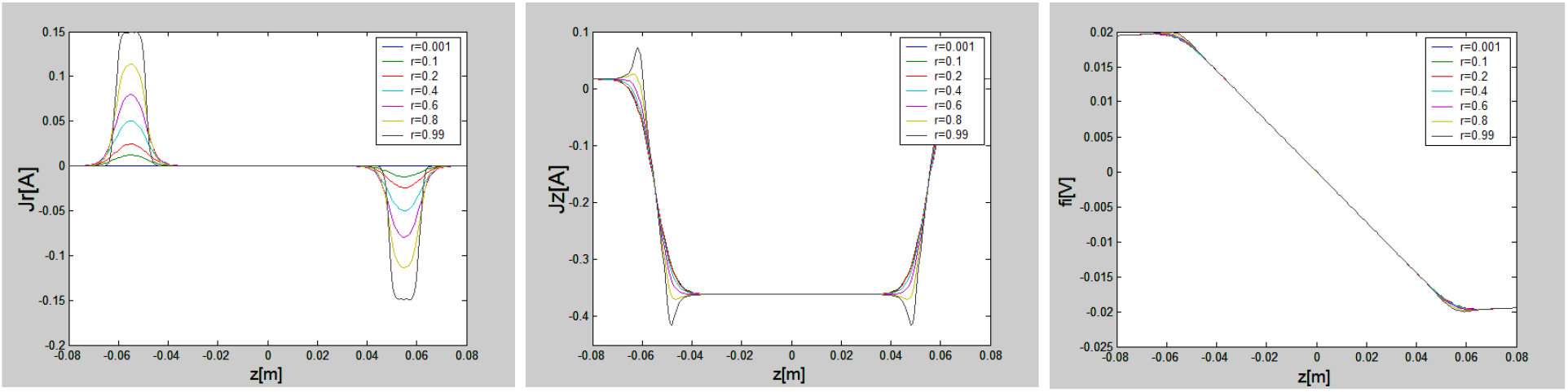
Dependences of currents and potential on *z* for various values of distances from the axis. The calculations were carried out for the selected parameters: *R*_0_ = 1 cm, *l*_0_ = 10 cm, *z*_1_ = 5 cm, *z*_2_ = 5.25 cm, *z*_3_ = 0.5 cm, *z*_4_ = 0.575 cm, *z*_5_ = 6 cm, *s* = 5 cm^−1^,*J*_0_ = 0.15 A.

## IV. Discussion

According to the results of the calculations for the considered example, the radial current (Fig. 3(A)) near the axis of the external dielectric cylinder tends to zero, and the potential (Fig. 3(B)) is practically independent of the radius. Thus, the change in potential across the membranes of the nodes of Ranvier (Fig. 1(B)) can be considered equal to the change in potential on the axis of the considered dielectric cylinder. Therefore, in accordance with the results of the calculations, the potential on the membrane of the nodes of Ranvier in the vicinity of the longitudinal coordinate corresponding to the left electrode becomes approximately 20 mV higher than the resting potential, and on the right one, 20 mV lower.

Consider the case of action potential propagation along the fiber from right to left. It was shown in [26, 27] that when an action potential is excited in the n^th^ node of Ranvier, the induced potential in adjacent (non-excited) nodes of Ranvier depends on the length of the myelinated segments. If the myelinated areas are long enough, then the action potential excitation threshold of 40–45 mV will be only at the (n + 1)^st^ node of Ranvier, as shown schematically in Figure 4.

**Figure 4.**
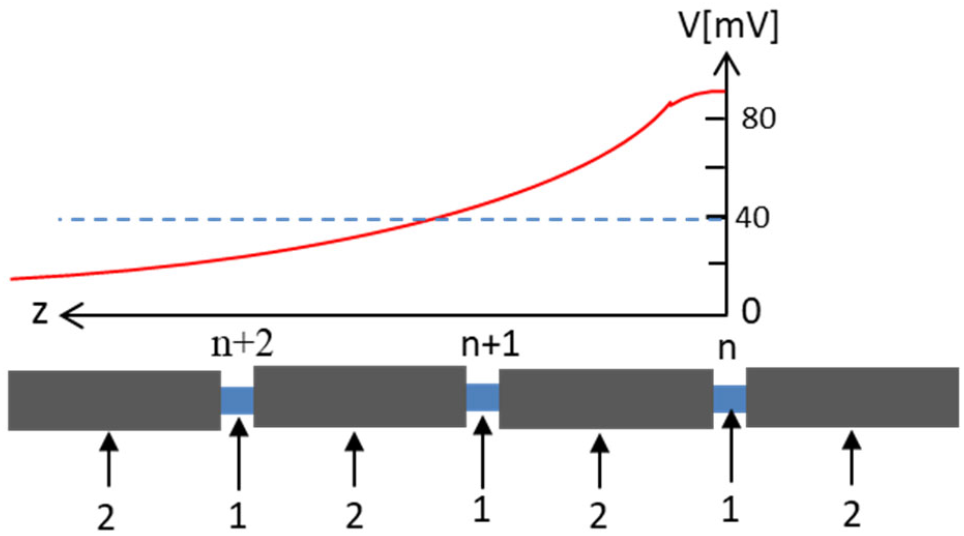
Schematic representation of an action potential propagating from left to right in a myelinated fiber. 1. Nodes of Ranvier; 2. Myelinated areas. The moment of time is shown when the action potential in the n^th^ node of Ranvier reached its maximum, in (n+1)^st^ the potential reached the threshold value, and in (n+2)^nd^ the potential did not reach the threshold excitation. The dotted line shows the value of the action potential corresponding to the threshold of excitation at the nodes of Ranvier.

According to the calculation results shown in Figure 3(C), the potential in the nodes of Ranvier near the right electrode is lowered by ∼20 mV relative to the resting potential, so the excitation threshold in them increased from 40–45mV to 60–65mV. Therefore, the propagation of the action potential near the right electrode is blocked. Obviously, an action potential propagating from left to right will also be blocked at the right electrode.

Let us now consider the case when the myelinated fiber is not on the cylinder axis. From the graphs shown in Figure 3, it can be seen that when moving to the right from the right electrode at a distance equal toelectrode size 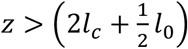, the current components *J* and *J* are close to 0, regardless of *r*, while the potential *φ* remains practically constant and independent of *r*. Therefore, the blocking of the action potential propagating from right to left can be expected to occur regardless of the radial position of the myelinated fiber. The same reasoning is also valid in the case of the propagation of the action potential from left to right: It will also be blocked behind the right electrode regardless of the radial position of the melein fiber.

Let us note a significant difference between the initiation of the action potential considered in [19] and the opposite task, blocking the action potential, presented in this work. In [19], a nerve fiber with axons located perpendicular to the currents induced in a conducting fluid was considered, and this, as shown in [19], greatly changes the structure of the currents near the axon. In this paper, we consider the case when mainly longitudinal currents are induced, and the radial current is close to zero. Therefore, it does not affect the potential and its distribution near the nerve fiber since the fiber is charged only by currents perpendicular to it. For the case of a fiber that is not on the axis of the considered dielectric cylinder with ring electrodes, all the considerations given above remain applicable.

## V. Conclusions

We have considered one of the possible approaches to anesthesia without the use of anesthetics. It has been shown that with a sequence of the excitation of current pulses in the vicinity of the nerve fiber, it is possible to block the propagation of the action potential, that is, a reversible loss of sensitivity without the use of anesthetics. However, it will be possible to draw final conclusions and talk about the application of the considered approach only after the experimental verification of the theory.

## Acknowledgments

The authors are grateful to E. Cohen and M. Keidar for valuable discussions.

